# DNALongBench: A Benchmark Suite for Long-Range DNA Prediction Tasks

**DOI:** 10.1101/2025.01.06.631595

**Authors:** Wenduo Cheng, Zhenqiao Song, Yang Zhang, Shike Wang, Danqing Wang, Muyu Yang, Lei Li, Jian Ma

## Abstract

Modeling long-range DNA dependencies is crucial for understanding genome structure and function across a wide range of biological contexts. However, effectively capturing these extensive dependencies, which may span millions of base pairs in tasks such as three-dimensional (3D) chromatin folding prediction, remains a significant challenge. Furthermore, a comprehensive benchmark suite for evaluating tasks that rely on long-range dependencies is notably absent. To address this gap, we introduce DNALongBench, a benchmark dataset encompassing five important genomics tasks that consider long-range dependencies up to 1 million base pairs: enhancer-target gene interaction, expression quantitative trait loci, 3D genome organization, regulatory sequence activity, and transcription initiation signals. To comprehensively assess DNALongBench, we evaluate the performance of five methods: a task-specific expert model, a convolutional neural network (CNN)-based model, and three fine-tuned DNA foundation models – HyenaDNA, Caduceus-Ph, and Caduceus-PS. We envision DNALongBench as a standardized resource with the potential to facilitate comprehensive comparisons and rigorous evaluations of emerging DNA sequence-based deep learning models that account for long-range dependencies.

## Introduction

Genomic DNA sequences are the blueprint of life, guiding the development of cellular complexities. Although protein-coding DNA sequences are responsible for diverse biochemical functions within organisms, most eukaryotic genomes predominantly consist of non-coding sequences interspersed with protein-coding regions. These non-coding sequences contain a variety of gene regulatory elements, such as promoters, enhancers, non-coding RNAs, and other functional elements, which orchestrate when and where genes are activated or silenced. Over the past two decades, large-scale functional genomic projects, such as the ENCODE project [1], have cataloged extensive collections of putative non-coding regulatory elements in the human genome. However, our understanding of how these elements regulate gene expression remains limited. A critical challenge lies in the fact that genomes are dynamically folded into multi-scale 3D structures within the cell nucleus, resulting in widespread physical DNA-DNA interactions, even between regions located megabases apart [2–4]. Effectively determining which of these interactions are functionally relevant to cellular processes across diverse biological contexts requires significant experimental effort.

To address this challenge, the increasing availability of genomic data, such as ChIP-seq [5], ATAC-seq [6], Hi-C, and its derivatives [7], has spurred the development of supervised deep learning methods that show great promise in systematically delineating sequence-to-function relationships. For example, convolutional neural networks (CNNs) and transformer-based methods have demonstrated their effectiveness in characterizing regulatory elements [8–11], predicting spatial proximity between genomic loci [12, 13], and predicting gene expressions from local sequence contexts [14]. Despite these advances, capturing dependencies across very long distal DNA elements remains a major computational challenge due to the scarcity of experimental data and the difficulty in modeling long-range sequence dependencies [15].

Recently, large language models (LLMs) have revolutionized the field of natural language processing (NLP), demonstrating remarkable capabilities across a wide spectrum of applications [16–19]. These models first leverage self-supervised learning techniques to learn intricate patterns from vast amount of unlabeled text data, followed by fine-tuning tailored to specific tasks. Recognizing structural similarities between DNA sequences and natural language [20], several DNA foundation models have emerged [21–23]. However, the utility of these models in addressing meaningful biological questions remains a topic of debate, leaving a critical question unsolved: *Could foundation models pre-trained on genomic DNA sequences offer a new paradigm shift in understanding the interactions between regulatory elements and genes?* Answering this question requires robust benchmark datasets to evaluate their performance, identify limitations, and drive future improvements. However, most foundation models pre-trained on genomic DNA sequences have so far been evaluated only on prediction tasks involving sequences up to a few thousand base pairs, such as identifying regulatory elements and predicting gene expression [24–28]. The potential of DNA LLMs to capture long-range interactions between DNA sequences in diverse biological contexts has not been well evaluated.

Here, we introduce DNALongBench, the largest collection to date of realistic and biologically meaningful genomic DNA prediction tasks that require long-range sequence input and involve long-range dependencies. DNALongBench comprises five different tasks and datasets, each addressing critical aspects of studying gene regulation across various length scales. The contributions of DNALongBench are threefold:

- We introduce DNALongBench, a benchmark for long-range DNA prediction tasks spanning up to 1 million base pairs (bp) across five distinct tasks. To the best of our knowledge, DNALongBench is the most comprehensive benchmark designed specifically for long-range DNA prediction tasks available to date.
- We evaluate the proposed DNALongBench using three representative models, demonstrating that while DNA foundation models can capture long-range dependencies to a certain extent, expert models consistently outperform DNA foundation models across all five tasks.
- The models exhibit varying performance across different tasks, highlighting the diverse challenges inherent in the DNALongBench prediction tasks and revealing the differing levels of difficulty associated with each task.

We envision DNALongBench as a valuable resource for evaluating foundation models trained on DNA sequences, with a particular focus on their ability to model long-range genomic interactions.

## Related Work

### Existing Benchmark datasets considering long-range DNA sequence

Benchmark datasets specifically designed to evaluate the capabilities of DNA foundation models in capturing long-range DNA dependencies remain underexplored. Most existing benchmarks for DNA foundation models primarily focus on short-range tasks (e.g., sequences spanning thousands of base pairs) and binary classification. To date, BEND [24] and the Genomics Long-range Benchmark (LRB) [28] are the only two existing benchmark datasets that include long-range genomic DNA prediction tasks. BEND comprises two long-range tasks: enhancer annotation and gene finding, both of which involve classifying regulatory elements. LRB, on the other hand, adapted all tasks from the Enformer [14] paper and curated three datasets focused on gene expression prediction and the effects of variants on gene expression. However, both BEND and LRB are limited in scope. They focus specifically on identifying regulatory elements or gene expression-related predictions and overlook other critical long-range DNA prediction tasks. For instance, neither benchmark includes structure-related predictive tasks requiring ultra-long sequences, such as contact map prediction and enhancer-target gene prediction. Furthermore, they lack base-pair-resolution regression tasks for quantitative assays. As a result, a comprehensive benchmark suite for evaluating a broader range of tasks reliant on long-range dependencies remains absent. We compare the scope of previous benchmarks for DNA prediction tasks with DNALongBench in **Table** 1.

**Table 1:** Comparison of existing benchmarks (Genomic Benchmarks [25], NT Benchmark [26], GUE [27], BEND [24], and LRB [28]) for DNA prediction tasks with DNALongBench. While recent benchmarks have been proposed for genomics, only BEND and LRB include tasks involving relatively long-range dependencies. In contrast, DNALongBench offers the most extensive range of long-range tasks, supporting sequences up to 1 million base pairs. Additionally, it features a greater variety of task types, longer input sequences, and evaluates the performance of three representative baseline models: supervised, expert, and DNA foundation models. The supervised baseline represents fully supervised models, such as lightweight CNNs, that do not involve pre-training.

| Benchmark Feature | Genomic Benchmarks | NT Benchmark | GUE | BEND | LRB | DNALONGBENCH |
| --- | --- | --- | --- | --- | --- | --- |
| Has Long-range Task | × | × | × | ✓ | ✓ | ✓ |
| Longest Input (bp) | 4,707 | 600 | 1,000 | 100k | 192k | 1M |
| Has Base-pair-resolution Regression Task | × | × | × | × | × | ✓ |
| Has two-dimensional Task | × | × | × | × | × | ✓ |
| Has Supervised Model Baseline | ✓ | × | × | ✓ | × | ✓ |
| Has Expert Model Baseline | × | × | × | ✓ | ✓ | ✓ |
| Has DNA Foundation Model Baseline | × | ✓ | ✓ | ✓ | ✓ | ✓ |

### Long-range DNA sequence modeling

Over the last decade, deep learning models for genomics have been dominated by CNNs [29]. While CNNs excel at extracting local features, their limited local receptive field constrains the flow of information between distant genomic elements [14]. To address this limitation, researchers have introduced dilated convolution and skipped connections to CNN architectures. For example, the Akita model [12] predicts chromatin contact maps (i.e., chromatin folding) from DNA sequences up to around 1 million base pairs by leveraging successive dilated convolutional layers and residual connections, enabling information flow across long distances.

Unlike CNNs, which require multiple successive layers to capture long-range dependencies, transformers use the attention mechanism, allowing each position in the sequence to directly attend to all other positions [30]. However, transformers are computational inefficient for long sequences, as the attention mechanism scales quadratically with sequence length [31]. This makes direct base-to-base attention across extremely long genomic sequences, spanning millions of base pairs, impractical [23, 31]. To address this, hybrid models combining convolutional layers for feature extraction with transformer modules have been proposed. The Enformer model [14], for instance, integrates convolutional layers with transformers to predict epigenetic and transcriptional features across long DNA sequences up to 200k bases in length.

Recently, DNA foundation models have emerged as an active area of research [14, 22, 23, 26, 27, 32, 33]. These models, pre-trained on large-scale DNA sequences, show promise across various downstream genomics tasks [22, 26, 27]. However, transformer-based DNA foundation models typically have relatively short context lengths (up to 4k tokens) due to the computational constraints of the self-attention mechanism [22, 26]. To address these limitations, researchers are exploring new transformer variants [34] and alternative architectures beyond transformers [22, 23, 31]. For instance, HyenaDNA [22] is a non-transformer-based DNA foundation model that uses implicit convolutions, allowing for long context lengths up to 1 million base pairs. It has demonstrated promising performance in long-range species classification tasks, although the practical applications of this problem remain poorly defined. Similarly, Caduceus [23] is a bi-directional equivalent long-range DNA foundation model built on Mamba blocks [31]. In this study, we selected HyenaDNA and Caduceus as part of the DNA foundation model baseline methods for evaluation in DNALongBench, as they are specifically designed for long-range DNA prediction tasks.

### Proposed Dataset: DNALongBench

The selection of suitable long-range DNA prediction tasks for DNALongBench is crucial to ensure diversity, comprehensiveness, and rigor. To achieve this, we established the following criteria to guide our task selection process:

#### Biological Significance

Tasks should be realistic and biologically significant, addressing genomics problems important for understanding genome structure and function.

#### Long-range Dependencies

Tasks should require modeling long input contexts, spanning hundreds of kilobase (kb) pairs or more.

#### Task Difficulty

Tasks should pose significant challenges to current models.

#### Task Diversity

Tasks should be as diverse as possible, spanning various length scales and including different task types, such as classification or regression. This diversity also encompasses task dimensionality (1D or 2D) and granularity (binned, nucleotide-wide, or sequence-wide).

As a result, we selected five long-range DNA prediction tasks, each covering various aspects of important regulatory elements and biological processes within a cell, as illustrated in **Fig**. 1. An overview of our dataset is presented in **Table** 2. The input sequences for all tasks are provided in BED format, which lists the genome coordinates of the sequences. This format allows for flexible adjustment of the flanking context without requiring reprocessing. The selected tasks are described in detail in the following sections.

**Figure 1:**
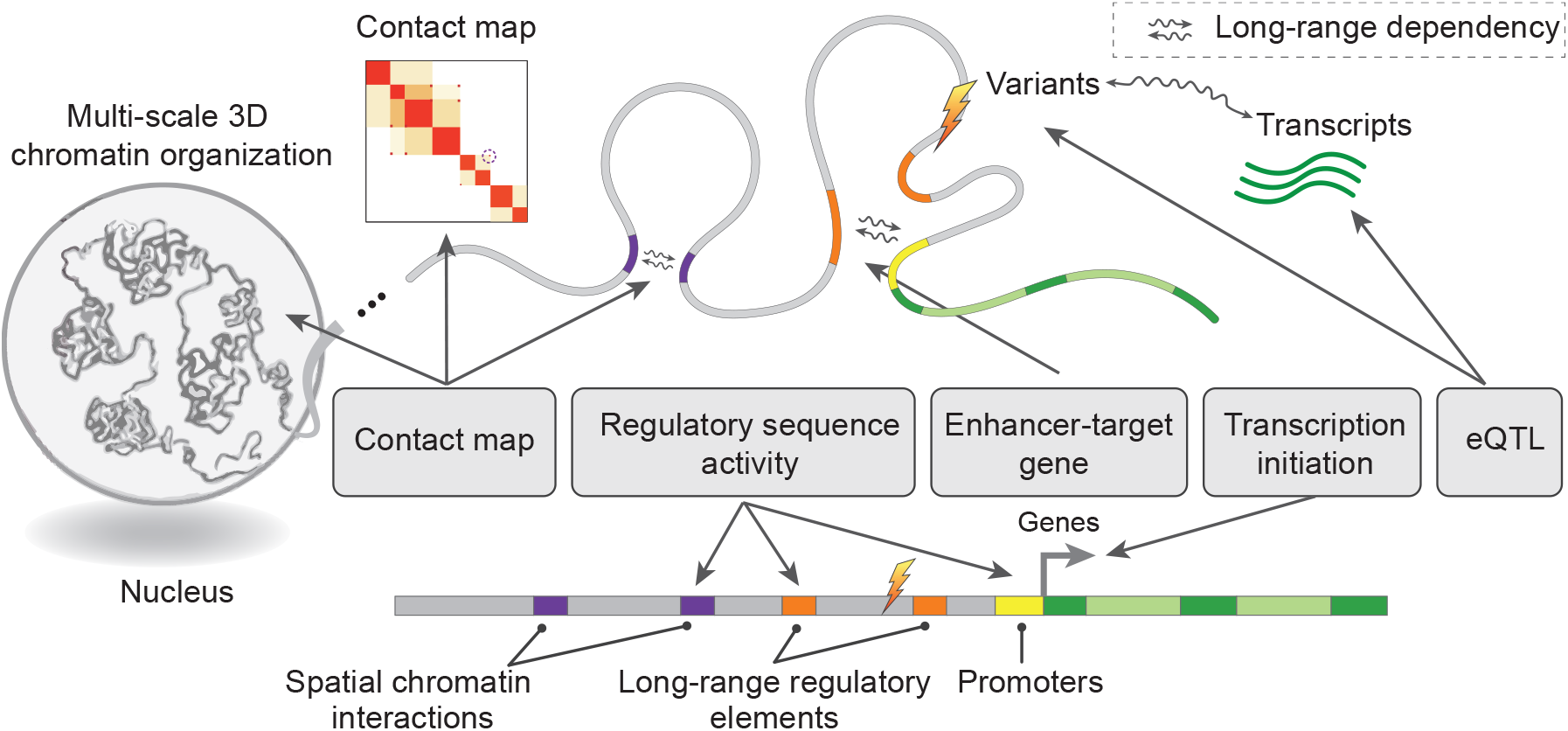
Illustration of the different categories of downstream tasks included in DNALongBench.

**Table 2:** Overview of the tasks included in DNALongBench. “1D” and “2D” denote one-dimensional and two-dimensional, respectively. Nucleotide-wise tasks involve predicting a sequence of labels, each corresponding to individual nucleotides in the input. Sequence-wise tasks require classifying the entire input sequence. In binned tasks, multiple nucleotides are grouped into bins and share a common label. *: The data for this task consists of sequences sampled from whole genomes, with 100,000 being the number of samples used for training our baselines. AUROC: Area Under the Receiver Operating Characteristic curve. PCC: Pearson correlation coefficient. SCC: stratum-adjusted correlation coefficient.

| LR Tasks | LR Type | Input Length | Output Shape | # Samples | Metric |
| --- | --- | --- | --- | --- | --- |
| Enhancer-target Gene | Binary Classification | 450,000 | 1 | 2602 | AUROC |
| eQTL | Binary Classification | 450,000 | 1 | 31282 | AUROC |
| Contact Map | Binned (2,048bp) 2D Regression | 1,048,576 | 99681 | 7840 | SCC&PCC |
| Regulatory Sequence Activity | Binned (128bp) 1D Regression | 196,608 | Human: (896, 5313)<br>Mouse: (896, 1643) | Human: 38171<br>Mouse: 33521 | PCC |
| Transcription Initiation Signal | Nucleotide-wise 1D Regression | 100,000 | (100000, 10) | 100000* | PCC |

### Enhancer-Target Gene Prediction

In eukaryotic cells, enhancers play a key role in gene regulation by forming enhancer-promoter interactions that activate the transcription of target genes, even those located up to several megabase pairs away [35]. However, the detailed mechanism by which sequence information encodes enhancer-promoter interactions remains poorly understood. Predictive methods that incorporate the full sequence information between enhancers and promoters as input could not only improve the prediction performance but also help identify the sequence determinants driving these interactions. To this end, we formulated a task to predict true enhancer-promoter interactions from a list of putative candidates based on the DNA sequence.

We collected experimentally verified enhancer-promoter interactions in K562 cells from [36–38]. Using CRISPRi-FlowFISH technique, the authors perturbed thousands of candidate sequences, quantified their effects on gene expression, and identified both positive and negative enhancer-promoter interactions. We filtered this data by retaining enhancer-promoter pair candidates within 450 kb of the gene transcription start site (TSS) and applied additional filtering criteria. Model performance was evaluated using AUROC. We compared models relying solely on sequence information with the expert model, activity-by-contact (ABC) model [36], which incorporates DNase-seq, H3K27ac ChIP-seq data, and a Hi-C matrix to prioritize true enhancer-promoter interactions. It should be noted that the ABC model inherent advantages over sequence-only models due to its more comprehensive input data types. The primary motivation here is to compare sequence-only models and understand their strengths and limitations.

### 3D Chromatin Contact Map Prediction

Chromosomes are folded in a well-organized manner within the cell nucleus, affecting various critical cellular functions such as gene transcription and DNA replication [39, 40]. Developing prediction models that connect 1D DNA sequences with 2D contact maps enables the identification of key sequence determinants of 3D chromatin folding, providing valuable insights into the underlying mechanisms of genome organization [4, 41]. We formulated a 3D chromatin contact map prediction task, defined as a 2D regression task to predict pairwise chromatin interactions between every pair of genomic loci within a given context window.

These contact frequencies are expressed as 2D contact maps derived from genomic mapping data such as Hi-C and Micro-C [4]. We used the processed data from Akita [12], which includes chromatin interaction data from five cell lines: HFF, H1-hESC, GM12878, IMR-90, and HCT116. To increase the number of cell types, we curated and processed additional Hi-C data for four cell lines: HAP1, Hela, HepG2, and K562, following the same data processing steps as in the Akita model. Each input sequence, spanning 1 million base pairs (Mbp), is divided into 512 genomic bins at a resolution of 2kb per bin. For the final prediction, 32 genomic bins are cropped from each side, resulting in a contact map of 448*×*448. Since the contact map is symmetric, predictions are made only for the upper triangular region, with a diagonal offset of 2. The human genome was split into non-overlapping virtual contigs and randomly assigned to training, validation, and testing sets with an 8:1:1 ratio. The dataset contains 7,008 training sequences, 419 validation sequences, and 413 test sequences. Model performance on the held-out test set was evaluated using the Stratum-Adjusted Correlation Coefficient (SCC) and the Pearson Correlation Coefficient (PCC).

### Regulatory Sequence Activity Prediction

Cell type-specific regulatory activities are encoded by the compositions and interactions of functional DNA segments, such as promoters, enhancers, and insulators, which can regulate genes from distant genomic locations. Predicting functional signals directly from DNA sequences over large genomic distances could help identify distal regulatory elements and uncover key sequence features enabling long-range gene regulation. For this task, we compiled human and mouse genomic tracks from the Enformer paper [14]. The goal of this task is to predict thousands of epigenomic profiles directly from DNA sequence spanning up to 100kb. We formulated the task as a multitask regression problem aimed at predicting epigenetic and transcriptional signals from long DNA sequences alone.

The dataset includes experimentally determined regulatory activity signal tracks and corresponding DNA sequences from human and mouse genomes. Each input DNA sequence spans 196,608 bp, centered on the TSS of protein-coding genes. Each input sequence consists of a core region and flanking regions. The core sequence is 114,688 bp in length, corresponding to 896 bins at a resolution of 128 bp per bin. The target labels consist of 5,313 human tracks and 1,643 mouse tracks measuring epigenomic marks. The dataset contains 38,171 human sequences and 33,521 mouse sequences. For the human genome, the data is split into 34,021 training, 2,213 validation, and 1,937 test sequences. For the mouse genome, the dataset is split into 29,295 training, 2,209 validation, and 2,017 test sequences. Model performance was evaluated using Pearson correlation coefficient, calculated by comparing predicted and target signal tracks. Specifically, the Pearson correlation coefficient was computed for each sample across all positions and tracks, and the mean was taken across all samples in the test set.

### Expression Quantitative Trait Loci (eQTL) Prediction

Expression quantitative trait loci (eQTL) are nucleotide variants that affect the expression of one or more genes. Deep learning-based approaches for predicting gene expression from DNA sequences have gained increasing popularity. One practical application of these methods is the identification and interpretation of eQTLs, a traditionally labor-intensive and time-consuming process when relying on genome-wide association studies. We designed an eQTL prediction task to provide an efficient approach for evaluating eQTLs, where the goal is to predict whether a nucleotide variant modulates the expression of a target gene using DNA sequence alone.

We adapted the eQTL dataset used in Enformer [14]. Positive SNPs were identified using the statistical fine-mapping tool Susie [42]. The original dataset includes positive and matched negative variants across 48 tissues [14]. For this study, we selected the top nine tissues based on the number of variants. Within these tissues, eQTL-gene pairs were filtered to retain eQTL candidate loci within 450 kb of the gene TSS. Genes with fewer than two positive pairs, two negative pairs, or five combined pairs were excluded. The sequences between variants and promoters were extracted, extending 3 kb downstream of the gene TSS. To reduce bias caused by putative eQTLs within the interval between an eQTL candidate and the gene promoter pair, we masked the sequences of all variants within each variant-promoter pair. The dataset was randomly split into training, validation, and test sets using a stratified sampling approach with an 8:1:1 ratio. To ensure robustness, at least one positive and one negative pair were included in both the training and validation sets. Model performance was evaluated using AUROC.

### Transcription Initiation Signal Prediction

Promoters are specialized DNA sequences at the TSS of genes that support the assembly of transcription machinery and transcription initiation [43]. Each promoter exhibits a unique profile of transcription initiation signals, which may reflect the mechanisms underlying transcription initiation. Solving the machine-learning task of predicting these profiles from promoter sequences provides insights into sequence-based regulation of transcription initiation [44]. Using long sequences as input and improving the information flow between distal elements could enhance predictive accuracy of transcription initiation signal prediction. We include a task in DNALongBench aimed at predicting transcription initiation signal profile from DNA sequences. Specifically, the task predicts transcription initiation signals on both strands for five experimental techniques: FANTOM CAGE, ENCODE CAGE, ENCODE RAMPAGE, GRO-cap, PRO-cap [44]. Unlike the regulatory sequence activity prediction task, which predicts sequence coverage at 128 bp genomic bins, this task requires predictions at base-pair resolution, making it significantly more challenging.

We used processed labeled data from the Puffin work [44]. Predictions were generated for entire test chromosomes (chr8 and chr9) using a sliding window step size of 50 kb, with the center 50 kb of each 100 kb prediction being evaluated. Regions within 1 kb of unknown bases or within 25 kb of chromosome ends were excluded. Model performance was evaluated using Pearson’s correlation.

Additional details on data processing and data access are provided in the **Supplemental Information**.

### Experiments

In this section, we conduct a comprehensive performance comparison by evaluating three distinct types of models: a lightweight convolutional neural network, existing expert models that have demonstrated state-of-the-art results, and two types of recent DNA foundation models – HyeynaDNA [22] and Caduceus [23] – distinguished by their support to reverse complement DNA during the training process.

### Representative Models

We explore the performance of the following three types of models:

1. **CNN**: We evaluate the lightweight convolutional neural network [45], known for its simplicity and robust performance across various DNA-related tasks. Detailed model implementations for each task are provided in **Supplemental Information**.
2. **Expert Model**: We assess the current state-of-the-art models for each specific long-range DNA prediction task, collectively referred to as the expert model. Specifically, we use:
  - The Activity-by-Contact (ABC) model [36] for the enhancer-promoter interaction prediction.
  - Enformer [14] for the eQTL prediction and regulatory sequence activity prediction.
  - Akita [12] for contact map prediction.
  - Puffin-D [44] for transcription initiation signal prediction.
  Additional details about each expert model are provided in **Supplemental Information**.
3. **DNA Foundation Model**: We selected three long-range DNA foundation models – HyenaDNA (medium-450k) and Caduceus (Ph and PS) – for evaluation. Due to limited computing resources, we were unable to fine-tune HyenaDNA-large-1m and Evo (7B) [21]. The detailed fine-tuning strategies for each task are provided in the **Supplemental Information**.

### Benchmarking Results

The main results are reported in **Tables** 3, 4, and 5. Additional metrics are provided in **Table** S1.

**Table 3:** AUROC for enhancer-target gene prediction (ETGP) task and SCC scores for the contact map prediction (CMP) task. K562, HFF, H1hESC, GM12878, IMR90, and HCT116 represent different human cell types. The highest scores are highlighted in bold. “Avg” means the average score across different cell types. Notably, the Expert Model achieves the best performance on both ETGP and CMP tasks.

| Models | ETGP |  | CMP |  |  |  |  |
| --- | --- | --- | --- | --- | --- | --- | --- |
|  | K562 | HFF | H1hESC | GM12878 | IMR90 | HCT116 | Avg |
| Expert Model | **0.926** | **0.258** | **0.247** | **0.227** | **0.210** | **0.210** | **0.230** |
| CNN | 0.797 | 0.025 | 0.024 | 0.010 | 0.013 | 0.001 | 0.015 |
| HyenaDNA | 0.828 | 0.139 | 0.122 | 0.099 | 0.097 | 0.118 | 0.115 |
| Caduceus-Ph | 0.826 | 0.153 | 0.130 | 0.101 | 0.138 | 0.145 | 0.133 |
| Caduceus-PS | 0.821 | 0.142 | 0.123 | 0.097 | 0.132 | 0.139 | 0.127 |

**Table 4:**
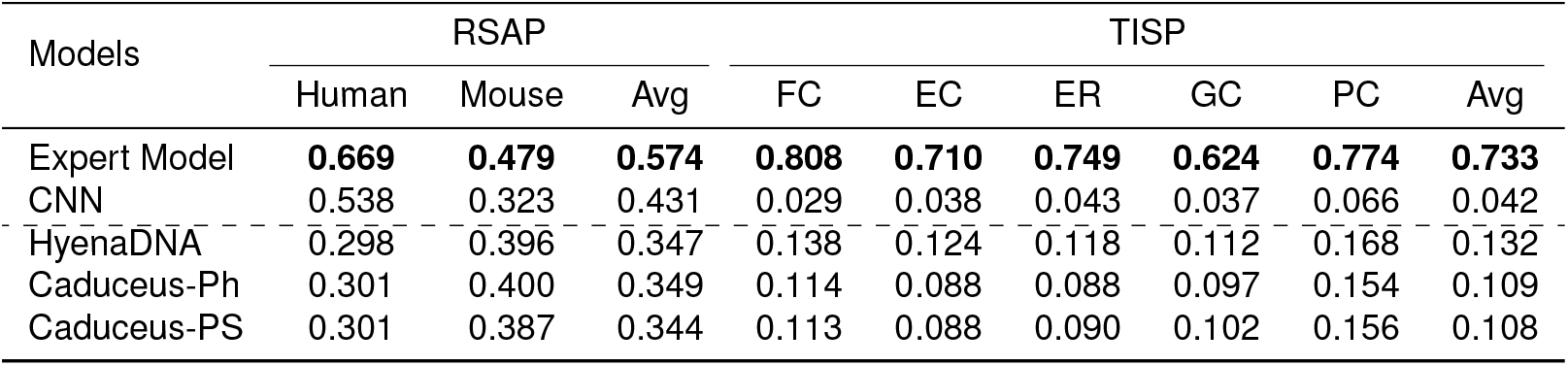
Pearson correlation scores for the regulatory sequence activity prediction (RSAP) task and the transcription initiation signal prediction (TISP) task. FANTOM CAGE (FC), ENCODE CAGE (EC), ENCODE RAMPAGE (ER), GRO-cap (GC), and PRO-cap (PC) represent the experimental techniques used in the TISP task. “Avg” means the average scores. The highest scores are highlighted in bold. Notably, the Expert Model significantly outperforms both the CNN and DNA foundation models.

**Table 5:** AUROC scores for the expression quantitative trait loci prediction (eQTLP) task across different cell types. The cell types are abbreviated as follows: CCF for Cells_Cultured_fibroblasts, WB for Whole_Blood, SNSES for Skin_Not_Sun_Exposed_Suprapubic, SSELL for Skin_Sun_Exposed_Lower_leg, MS for Muscle Skeletal, NT for Nerve Tibial, AT for Artery_Tibial, and AS for Adipose_Subcutaneous. The highest scores are highlighted in bold. “Avg” represents the average score across all cell types. Notably, the Expert Model achieves the best performance across all cell types.

| Models | eQTL |  |  |  |  |  |  |  |  |  |
| --- | --- | --- | --- | --- | --- | --- | --- | --- | --- | --- |
|  | CCF | WB | Thyroid | SNSSES | SSELL | MS | NT | AT | AS | Avg |
| Expert Model | **0.639** | **0.689** | **0.612** | **0.710** | **0.700** | **0.621** | **0.683** | **0.741** | **0.736** | **0.681** |
| CNN | 0.547 | 0.577 | 0.487 | 0.499 | 0.499 | 0.502 | 0.516 | 0.576 | 0.551 | 0.528 |
| HyenaDNA | 0.584 | 0.512 | 0.529 | 0.471 | 0.544 | 0.487 | 0.511 | 0.479 | 0.513 | 0.514 |
| Caduceus-Ph | 0.597 | 0.594 | 0.527 | 0.586 | 0.574 | 0.538 | 0.588 | 0.547 | 0.541 | 0.565 |
| Caduceus-PS | 0.549 | 0.542 | 0.547 | 0.529 | 0.541 | 0.523 | 0.552 | 0.536 | 0.519 | 0.537 |

#### The Expert Model achieves the highest scores on all tasks

Specifically, the Expert Model achieves an average score of 0.733 on the transcription initiation signal prediction task (TISP), significantly surpassing CNN’s 0.042, HyenaDNA’s 0.132, Caduceus-Ph’s 0.109, and Caduceus-PS’s 0.108. This disparity may stem from the challenge posed by multi-channel regression on long DNA contexts, which make fine-tuning of DNA foundation models less stable and less capable of capturing sparse real-valued signals. In the remaining four tasks, the Expert Model still outperforms both CNN and DNA foundation models, though the margin is less pronounced. For instance, Caduceus-Ph’s performance on the contact map prediction task (CMP) is only slightly lower than the Expert Model and much better than CNN. Overall, these observations confirm the Expert Model’s superior ability to capture long-range dependencies, a capability where CNN falls short and DNA foundation models demonstrate good performance in certain tasks.

#### The regulatory sequence activity prediction presents greater challenges

In contrast to the other four tasks, where the Expert Model or DNA foundation models achieve reasonable performance, the regulatory sequence activity prediction task proves significantly more difficult. The highest average Pearson correlation score achieved in this task is 0.574 by the Expert Model (Enformer), indicating only a moderate positive correlation. This result underscores the challenge of capturing long-range dependencies in regulatory sequence activity prediction and highlights the varying levels of task difficulty in DNALongBench.

### Analysis: Diving Deep into DNALongBench

In this section, we provide further analysis to gain insights into how long-range dependencies are captured in our proposed DNALongBench.

#### Case Study: Can long-range dependency be captured?

To intuitively demonstrate the presence of extensive long-range dependencies exist across millions of base pairs and their capture by machine learning methods, we present two examples in **Fig**. 2 and more examples in **Fig**. S1. Specifically, in **Fig**. 2a-b, we visualize the contact maps predicted by HyenaDNA, Caduceus-Ph, and the Expert Model (Akita), alongside the ground truth contact maps for two genomic regions spanning around 400 kb. From these contact maps, we observe the presence of large-scale domains (e.g., blocks in the contact map) and long-range interactions (e.g., dots in the contact map) spanning over 300 kb. Notably, the contact maps predicted by Akita align more closely with the ground truth, confirming its superior ability to capture long-range interactions. In contrast, the DNA foundation models show limited capacity for predicting domain structures. This is particularly evident in **Fig**. 2b, where only Akita accurately predicts the three blocks. These examples highlight DNALongBench’s valuable in evaluating models for capturing long-range genome structure and function and provide a foundation for future developments in DNA foundation models.

**Figure 2:**
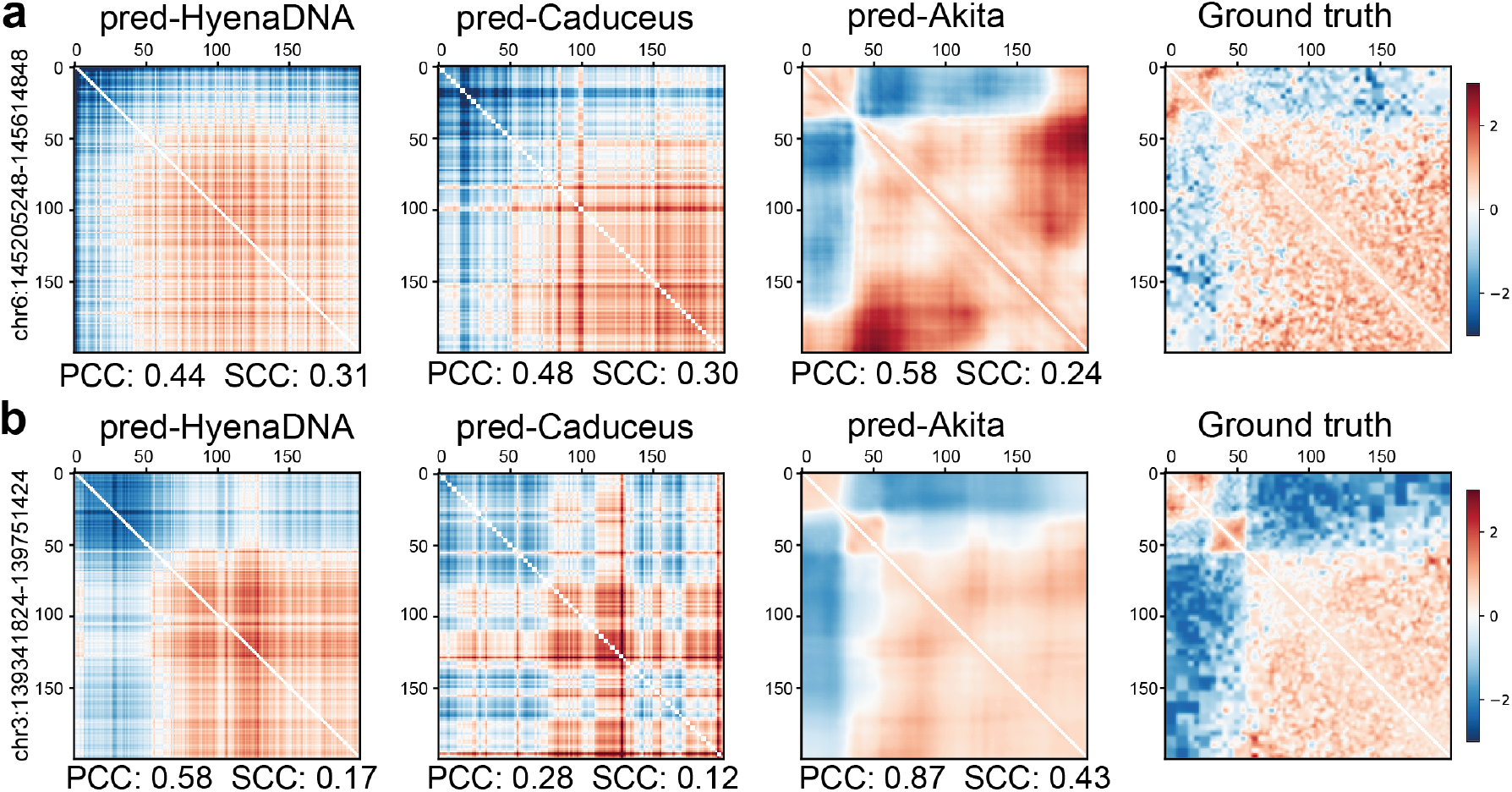
Comparisons of HyenaDNA, Caduceus (Ph), and the Expert Model (Akita) on the 2D contact map prediction task across 409,600 bp with a bin size of 2,048 bp. The columns show contact maps predicted by HyenaDNA, Caduceus, Akita model, alongside the ground truth contact map for two genomic regions: **(a)** chr6:145,205,248-145,614,848 and **(b)** chr3:139,341,824-139,751,424. Colors represent the intensity of contact frequency between paired loci. Pearson correlation coefficient (PCC) and stratum-adjusted correlation coefficient (SCC) metrics are shown beneath each contact map to indicate prediction performance relative to the ground truth.

#### Base pair-resolution prediction of transcription initiation signal

We visualized transcription initiation signals predicted by different models for one of the test chromosomes, chr8 (**Fig**. 3). Predictions from the Expert model, Puffin-D, closely align with the ground truth, accurately capturing peaks in transcription initiation signal intensity across both large and small genomic regions. By constrast, DNA foundation models tend to underpredict signal intensities or miss certain peaks. In the zoomed-in view (right side of the figure), Puffin-D continues to align well with the ground truth, demonstrating strong performance even at high resolution. In contrast, the DNA foundation models show broader, less precise signals. These findings suggest that base pair-resolution regression tasks remain challenging for DNA foundation models.

**Figure 3:**
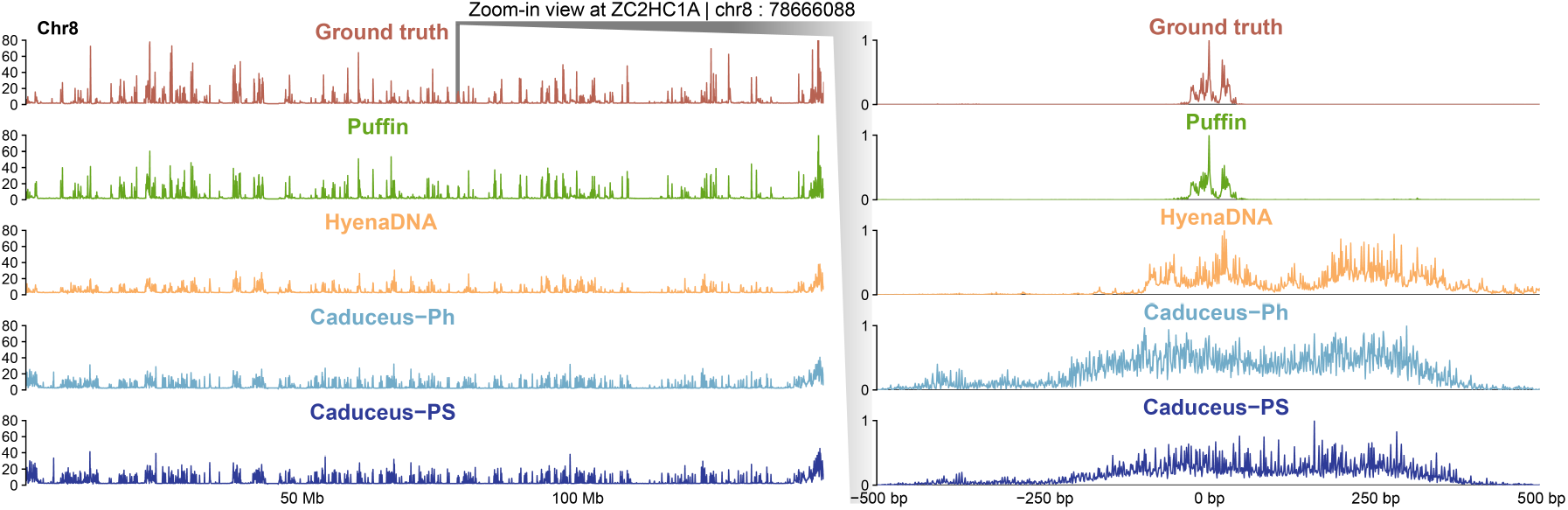
Comparisons of HyenaDNA, Caduceus_Ph, Caduceus_PS, and Expert Model (Puffin-D) on the transcription initiation signal prediction task of chromosome 8. The genomic track on the left displays the ground truth signals (top) alongside predictions from Puffin-D, HyenaDNA, and the two Caduceus models. The x-axis represents genomic coordinates, while the y-axis indicates signal density. A zoomed-in view of a 1,000 bp region centered at the TSS of the gene *ZC2HC1A* is shown on the right.

## Conclusion

In this paper, we introduce DNALongBench, a benchmark suite comprising five important genomics tasks involving long-range dependencies: enhancer-target gene interaction, eQTL, 3D genome organization, regulatory sequence activity, and transcription initiation signal. We evaluated three baseline methods: a task-specific expert model, a fully supervised CNN-based model, and three fine-tuned DNA foundation models, HyenaDNA, Caduceus-Ph and Caduceus-PS. The benchmarking results consistently demonstrated that expert models achieved the highest scores across all tasks. Additionally, our analysis revealed that long-range dependencies could be captured across hundreds of thousands of base pairs, underscoring the importance of context length for downstream performance. However, the results also highlight that current DNA foundation models are less effective than expert models in capturing long-range dependencies. Nevertheless, we believe that DNALongBench will serve as a valuable resource for facilitating comprehensive comparisons and rigorous evaluations of emerging DNA sequence-based deep learning models that account for long-range dependencies.

One limitation of this study is the exclusion of transformer-based DNA foundation models, such as DNABERT-1, DNABERT-2, and Nucleotide Transformer, due to the computational challenges posed by training them on long-range tasks. The quadratic cost of the self-attention mechanism renders such tasks infeasible for these models. Exploring strategies to extend the context length of transformer-based models and effectively fine-tune them for long-range tasks remains an important avenue for future research, albeit beyond the scope of this study.

## Acknowledgment

This work was supported, in part, by National Institutes of Health Common Fund 4D Nucleome Program grant UM1HG011593 (J.M.); National Institutes of Health Common Fund Cellular Senescence Network Program grant UH3CA268202 (J.M.); and National Institutes of Health grants R01HG007352 (J.M.), R01HG012303 (J.M.), U24HG012070 (J.M.), and R21DA061481 (J.M.). J.M. also received support from a Guggenheim Fellowship awarded by the John Simon Guggenheim Memorial Foundation, a Google Research Award, and a Single-Cell Biology Data Insights Award from the Chan Zuckerberg Initiative. L.L. is supported by an NEC Faculty Research Award and the Neocortex Award from the Pittsburgh Supercomputing Center.

## Code Availability

The source code can be accessed at https://github.com/wenduocheng/DNALongBench.

## Data Availability

Datasets included in DNALongBench are available at https://tinyurl.com/dnalongbench

## Author Contributions

Conceptualization, W.C., Z.S., Y.Z., L.L., J.M.; Methodology, W.C., Z.S., Y.Z.; Software, W.C., Z.S., Y.Z., S.W., D.W., M.Y.; Investigation, W.C., Z.S., Y.Z., S.W., D.W., M.Y., L.L., J.M.; Writing, W.C, Z.S., Y.Z., J.M.; Funding Acquisition, L.L., J.M.

## Competing Interests

The authors declare no competing interests.

## Notes

### Competing Interest Statement

The authors have declared no competing interest.

