## Supplemental Information for "DNALongBench: A Benchmark Suite for Long-Range DNA Prediction Tasks"

#### A Dataset Documentation and Intended Uses

We document datasets of this study following the Datasheets for Datasets framework [1], discussing motivation, composition, collection process, preprocessing, and uses.

##### A.1 Enhancer-Target Gene Prediction

**Motivation**. This dataset is designed to explore the predictive power of DNA sequences alone in identifying the correct target genes of an enhancer. It can be used to evaluate DNA large language models for their ability to capture long-range dependency in regulatory elements.

**Composition**. This dataset contains multiple DNA sequences represented by nucleotide letters A, C, G, T, and N, where N indicates an unknown nucleotide or serves as padding. The sequences represent the DNA sequence between enhancers and their putative target genes. Genes are always located on the 5' end of the sequence, while enhancers are located on the 3' end. Sequences are fixed at 450,000 bp, with N as the padding added to the 3' end when the enhancer-gene distance is less than 450,000 bp. Each sequence is labeled as binary, indicating whether the enhancer regulates the target gene. The dataset contains 2,602 samples split into 2,066 training sequences, 266 validation sequences, and 270 test sequences. Stratified sampling ensures that each gene has at least one positive and one negative pair in both training and validation sets. The dataset includes tested enhancers for 24 genes.

**Collection Process**. The original enhancer-gene pairs were reported in [2] using the CRISPRi-FlowFISH method to experimentally determine regulatory relationship. For each gene, dozens of enhancer candidates were tested for their abilities to regulate gene expression. Enhancers significantly altering gene expression (FDR $<0.05$) were considered positive, while the remaining enhancers were labeled negative for that gene.

**Preprocessing**. Data was curated from [2–4], and sequences between enhancers and promoters were extracted using the hg19 reference genome. Enhancers located more than 450,000 bp from the TSS were excluded, and genes with fewer than two positive pairs, two negative pairs, or five combined pairs were removed. Sequence were masked between enhancers and promoters, extending 500 bp upstream of the enhancer and 3kb downstream of the gene TSS. To remove bias from potential enhancers located within the interval between an enhancer and promoter pair, the sequence of all intervening enhancers were masked. The entire dataset was randomly split into training, validation, and test sets using a stratified sampling approach with an 8:1:1 ratio. A restriction was applied to ensure that each split contained at least one positive and one negative pair. Since the sequences surrounding the enhancers and the TSS may contribute to gene regulation, we included an additional 3,000 bp of sequence downstream of the TSS and 500 bp upstream of each enhancer candidate. Additionally, the DNA sequence of other enhancers tested in the referenced study were masked to remove potential biases caused by these additional enhancers. The letter 'N' was added to the 3' end of each sequence to ensure a uniform length of 450,000 bp across all samples.

**Uses**. The authors of [2] used this dataset to evaluate the performance of their proposed activity-by-contact (ABC) model.

##### A.2 Contact Map Prediction

**Motivation**. The Contact Map Prediction dataset was created by the authors of Akita [5] to explore the connection between DNA sequences and 3D chromatin folding. Given a DNA sequence, a model predicts the interactions between each pair of genomic bins within the sequence.

**Composition**. The dataset contains human genome sequences and chromatin pairwise contact maps. Each sample consists of an input DNA sequence of length 1,048,576 bp, divided into 2,048-bp genomic regions, resulting in 512 bins per sequence. The output includes pairwise interaction frequencies for the central 448 bins, represented as a 448×448 2D matrix. Since the contact map is symmetric, only the upper triangular region is used for prediction, with a diagonal offset of 2. The final output is a vector of length 99,681. The dataset includes 7,840 sequences, partitioned into 7,008 training sequences, 419 validation sequences, and 413 test sequences.

**Collection Process**. Human genome sequence were collected from the human reference genome assembly GRCh38. Contact maps were derived from publicly available Hi-C and Micro-C data. The dataset includes five cell types: HFF and H1-hESC [6], GM12878 and IMR-90 [7], and HCT116 [8]. Additional data for four other cell lines were collected from the 4DN Data Portal. The data were processed following the Akita approach. Raw interaction pairs were binned into 2,048 bp bins, and functions in cooltools were used for normalization, adaptively coarse-grain smoothing, and linear interpolation of missing bins, and convolution with a small 2D Gaussian filter (sigma=1, width=5).

**Preprocessing**. The pre-processing steps include adaptive coarse-graining, normalization to account for distance-dependent decreases, log transformation, value clipping to the range of (-2, 2), linear interpolation of missing values, and convolution with a small 2D Gaussian filter.

**Uses**. This dataset was used by Akita [5] to predict chromatin interactions from DNA sequences.

##### A.3 Regulatory Sequence Activity Prediction

**Motivation**. The Regulatory Sequence Activity Prediction dataset aims to study the effects of DNA sequences on various epigenetic and transcriptional signals. A model trained on this dataset can infer the regulatory impact of noncoding DNA on gene expression.

**Composition**. The dataset consists of DNA sequences, each 196,608 bp in length, including 38,171 human sequences and 33,521 mouse sequences. For each sequence, the prediction targets are genome-wide tracks representing 5,313 features for human and 1,643 features for mouse. These tracks are measured using ChIP-seq, DNase-seq, ATAC-seq, and CAGE.

**Collection Process**. The dataset was developed by the authors of Enformer [9], extending the Basenji2 dataset [10] by increasing the length of input DNA sequences from 131,072 bp to 196,608 bp.

**Preprocessing**. The dataset uses the same training, validation, and test splits provided in the Enformer study. No additional preprocessing steps were applied.

**Uses**. The dataset has been used to train the Enformer model [9].

#### A.4 eQTL Prediction

**Motivation**. This dataset explores the predictive power of using DNA sequences alone to identify expression quantitative trait loci (eQTL) from germline mutations.

**Composition**. The dataset contains nine sub-datasets representing putative eQTL from nine human tissues. Each sample consists of a pair of DNA sequences (reference and alternative), each 450,000 bp in length, spanning the region between the mutation and the TSS of the target gene. The only difference between the reference and alternative sequences is the nucleotide at the variant site. Genes are located on the 5' end of the sequence, and mutations on the 3' end. Padding with 'N' is applied to the 3' end for sequences shorter than 450,000 bp. Each sample is labeled as either positive (eQTL) or negative. Sample counts range from 2,181 to 4,919 across tissues, with a total of 31,282 samples.

**Collection Process**. The dataset was originally created by the authors of Enformer [9], who collected statistical fine-mapping variants from GTEx v8 using the SuSiE method [11].

**Preprocessing**. The top nine tissues were selected based on variant count. Variant-gene pairs were filtered to include variants within 450,000 bp of the gene TSS. Sequences spanning the region between variants and genes were extracted, with an additional 3 kb downstream of the TSS. To avoid biases, intervening variants within the sequence were masked. Positive variants were identified based on posterior inclusion probability (PIP) $>0.9$, while negative variants were matched based on PIP $<0.01$ and specific Z-score thresholds. Stratified sampling was used to split the dataset into training, validation, and test sets (80:10:10), ensuring each split included at least two positive and two negative variants per gene.

**Uses**. This dataset has been used to evaluate the Enformer model [9] in predicting the effects of variants on gene expression.

#### A.5 Transcription Initiation Signal Prediction

**Motivation**. The Transcription Initiation Signal Prediction dataset investigates how DNA sequences determine transcription initiation. The task is to predict transcription initiation signals at base-pair resolution from DNA sequences.

**Composition**. The dataset includes human genome sequences and transcription initiation signals generated by five experimental techniques: FANTOM CAGE, ENCODE CAGE, RAMPAGE, GRO-cap, and PRO-cap. Each sample consists of a 100,000 bp DNA sequence and a 10-channel label representing signal intensities for both forward and reverse strands across the five techniques. The training set includes 100,000 intervals sampled from all chromosomes except 8, 9, and 10, which are used for validation and testing.

**Collection Process**. The dataset was created by the authors of Puffin [12]. Genome sequences were sourced from the GRCh38 reference genome, and transcription initiation signals were obtained from FANTOM [13] and ENCODE [14] projects, RAMPAGE [15], GRO-cap [16], and PRO-cap [17].

**Preprocessing**. The same random seed as Puffin-D was used to sample training sequences. Sampling continued until 100,000 intervals were obtained for training.

**Uses**. This dataset has been used to train the Puffin model [12].

### B License

DNALONGBENCH is licensed under CC BY 4.0. A copy of the license is provided with the dataset. Users are required to cite the original sources when utilizing the data provided with DNALONGBENCH. The authors bear full responsibility in case of any rights violations.

### C Implementation Details

All experiments were conducted on single A6000 GPUs within cluster environments. All models were implemented using PyTorch [18].

#### C.1 CNN Implementation Details

**Enhancer-Target Gene Prediction**. The CNN model was trained with cross-entropy loss and a learning rate of 0.005. The batch size was set to 5, and the model was optimized using Adam. The architecture consists of three Conv1D layers with 128, 64, and 32 filters, respectively, followed by a fully connected layer for class probability prediction. Max pooling was applied before the fully connected layer. The kernel size for all convolutional layers was set to 3 with padding of 1.

**Contact Map Prediction**. The CNN model was trained using Adam with a learning rate of 0.005 for 30 epochs and a batch size of 16. The architecture, designed specifically for this 2D regression task, consists of the following components:

- 1D convolutional tower: three Conv1D layers.
- Bottleneck layer: A 1D convolutional layer with 64 filters and a kernel size of 1.
- 2D convolutional tower: The bottleneck output is unsqueezed and repeated to create a tensor of shape [batch size, 64, 512, 512]. Positional encoding is added, resulting in a tensor of shape [batch size, 65, 512, 512], which is passed through three 2D
- Cropping layer: The 2D output is cropped to remove 32 bins from each side, yielding a tensor of shape [batch size, 64, 448, 448].
- Transformation layer: The upper triangular portion of the 2D map (diagonal offset = 2) is extracted, resulting in a tensor of shape [batch size, 64, 99681]. A final 1D convolutional layer reduces the feature dimension to 1, producing the final output tensor of shape [batch size, 1, 99681].

**Regulatory Sequence Activity Prediction**. The CNN model was trained with Poisson loss, a learning rate of 0.001, and a batch size of 16. The architecture consists of five convolutional layers with filter sizes of 16, 64, 256, 1024, and 5313 for human, and 16, 64, 256, 1024, and 1643 for mouse. Kernel sizes were set to 25, 15, 15, 15, and 1, respectively. Batch normalization and max pooling were added between each pair of layers, with adaptive max pooling applied after the fourth layer to ensure the output sequence length matched the target length of 896.

**eQTL Prediction**. The CNN model was trained with cross-entropy loss and a learning rate of 0.005. The batch size was set to 5, and the model was optimized using Adam. The architecture mirrored that of the enhancer-target gene prediction task, consisting of three Conv1D layers with 128, 64, and 32 filters, followed by a fully connected layer for class probability prediction. Max pooling was applied before the fully connected layer, and the kernel size for all convolutional layers was set to 3 with padding of 1.

**Transcription Initiation Signal Prediction.** The CNN model was trained with MSE loss, a learning rate of 0.005, and a batch size of 16. The architecture includes three Conv1D layers with 16, 32, and 64 filters.

#### C.2 Expert Model Details

**Enhancer-Target Gene Prediction**. We applied the Activity-by-Contact (ABC) model to predict whether an enhancer candidate is a true enhancer. Specifically, the ABC score for each enhancer of its putative target genes, as provided by [2], was used to calculate the AUROC on the testing set.

**Contact Map Prediction**. We used the Akita model [5] as the expert model. Akita consists of two main components: a trunk that learns 1D representations of DNA sequences and a head that converts these representations into 2D contact map predictions.

The trunk is composed of 11 convolutional blocks followed by 8 dilated residual 1D convolutions with geometrically increasing dilation rates. Each convolution block performs 1D convolution with 96 filters of width 5, batch normalization, ReLU activation, and maximum pooling of width 2. At the end of the trunk, a bottleneck convolutional layer with 64 filters and a width of 1 is added, resulting in an output of shape [512 bins, 64 filters]. The head converts the 1D sequence representation ([512, 64]) into a 2D map ([512, 512, 64]) by averaging representations between every pair of genomic bins $i$ and $j$. Additionally, the distance between bins ($|i - j|$) is included as an extra positional feature. Six blocks of dilated 2D convolutions, with geometrically increasing dilation rates, are applied to learn the contact map. After each block, resymmetrization is performed by averaging the contact map with its transpose. A linear transformation is used to predict contact maps for all five cell types simultaneously. We evaluated the pre-trained Akita model rather than training it from scratch.

**eQTL Prediction**. Following the Enformer paper [9], we first used the pre-trained Enformer model to extract features from reference and alternative sequences. Variants were represented as prediction difference vectors, calculated by subtracting these two features and summing the differences across the sequence. A random forest classifier was then employed to predict whether a variant was positive or negative. The random forest model was implemented using scikit-learn with 100 trees, and the maximum number of features considered for the best split was set to $\log_2$ of the total number of features.

**Transcription Initiation Signal Prediction.** We used the Puffin-D model [12] as the expert model for this task. Puffin-D is a CNN-based model designed with two upward and downward passes, connected by residual connections. Its architecture includes two upward blocks, two downward blocks, and one output block. The upward blocks consist of strided convolutional layers followed by batch normalization. The downward blocks include upsampling, strided convolutional layers, and batch normalization. The final output block comprises 1D convolutional layers with a kernel size of 1, batch normalization, ReLU activation, and Softplus activation. Residual connections, similar to U-Net [19], are implemented between corresponding levels of the upward and downward passes.

**Regulatory Sequence Activity Prediction**. We evaluated the performance of the pre-trained Enformer model* instead of training it from scratch [9].

*https://huggingface.co/EleutherAI/enformer-official-rough

##### C.3 DNA Foundation Model Finetuning Details

**Enhancer-Target Gene Prediction.** The feature vector was calculated by averaging hidden representations across the entire DNA sequence. This vector was used to perform a binary classification task, with all model parameters fine-tuned for optimal performance.

**Contact Map Prediction.** Feature vectors were calculated for each base pair bin (bin size: 2,048 bp), resulting in L/2048 bins, where L is the original DNA sequence length. A two-layer multi-layer perceptron (MLP) was used to calculate the contact map scores by taking concatenated feature vectors from two bins as input. This resulted in a contact map matrix of dimensions [L/2048, L/2048]. The upper triangular portion of this matrix was extracted, and mean squared error was applied to train the model. All parameters were fine-tuned.

**Regulatory Sequence Activity Prediction.** Following the standard split from Enformer [9], separate models were trained for human and mouse. Using 128 bp as a bin size, this task was formulated as multi-regression, with outputs of [896, 5313] for human and [896, 1643] for mouse. Poisson loss was used, with a batch size of 32 and a maximum training step of 30,000.

**eQTL Prediction.** The dataset consisted of triples (<original sequence, variant sequence, binary label>), with labels indicating whether a variant effect was positive or negative. Hidden representations from the last layer of both sequences were averaged and concatenated. A binary classification layer was applied to predict whether the variant was positive. All parameters were fine-tuned.

**Transcription Initiation Signal Prediction.** This task was formulated as a 10-channel regression problem using pseudo-Poisson KL divergence as the loss function. A linear layer predicted logits for 10 signals at each position. Following Puffin [12], 100k positions were randomly sampled from the genome sequence during training. The batch size was 16, with a maximum training step of 25,000.

#### D Additional Metrics

To evaluate model performance in the Contact Map Prediction task, we used two key metrics: PCC (Pearson Correlation Coefficient) and SCC (Stratum Adjusted Correlation Coefficient). While PCC is commonly used for comparing contact matrices, it does not account for domain structures or distance dependence—key characteristics of contact maps. To address these limitations, SCC, introduced by [20], enables more fine-grained differentiation between contact maps. Thus, we included both metrics in our evaluations. **Table** S1 presents the benchmark results using PCC, and **Fig.** S1 provides additional examples comparing predictive models with ground truth.

#### E Ablation Study on Context Length

To validate the importance of long-range context dependencies for task performance, we evaluated models with varying input context lengths.

##### E.1 Contact Map Prediction

Using Caduceus-Ph as an example, we evaluated performance with input sizes of 409,600, 307,200, and 204,800 bps, corresponding to 200, 150, and 100 bins, respectively. SCC was used as the evaluation metric. Results, shown in **Table** S2, indicate a clear trend: performance declined as context length decreased.

##### E.2 Regulatory Sequence Activity Prediction

For the regulatory sequence activity prediction task, using HyenaDNA as an example, we evaluated performance with sequence lengths of 196k and 131k bps. Results, shown in **Table** S3, also demonstrate a decline in performance with reduced context length.

##### E.3 Enhancer-Target Gene Prediction

Rather than reducing input size, we preserved enhancer sequences and regions near gene promoters, as these are critical for interactions. To study the effect of context length, we shuffled the central portion of the input sequence (50% of the original length). Using Caduceus-Ph as an example, **Table** S4 shows a notable decline in performance with central shuffling, suggesting the intervening DNA sequence is vital for predicting enhancer-gene interactions, consistent with previous studies [21].

##### E.4 Expression Quantitative Trait Loci (eQTL) Prediction

For the eQTLP task, we observed a similar trend using Caduceus-Ph (**Table** S5). Central shuffling led to a performance decline, confirming that long sequence contexts are beneficial for better results.

##### E.5 Transcription Initiation Signal Prediction

For the TISP task, using the expert model Puffin-D, we evaluated performance on holdout chromosomes 8–10 with varying effective contexts. Original predictions focused on the central 50kb of each 100kb input sequence. To examine the effect of flanking regions, we shuffled sequences in the first and last 25kb regions. Results, shown in **Tables** S6 and S7, reveal a slight decrease in accuracy with shorter contexts.

#### F Results on Additional Enhancer-Target Gene Prediction Datasets

We collected two additional CRISPRi screening-based enhancer-target gene datasets to evaluate model performance. CRISPRi screening methods are among the most direct and accurate for determining enhancer regulation of target genes. We also evaluated an additional expert model, gABC, as its authors reported superior performance over ABC [22].

Results for the additional datasets are presented in **Tables** S8 and S9. These findings show that Caduceus-Ph and gABC achieve the highest performance on their respective datasets.

#### G Results on Additional Contact Map Prediction Prediction Datasets

We curated and processed four additional datasets for the contact map prediction task by downloading Hi-C data for four cell types from the 4DN data portal [23, 24] and processing it using the Akita methodology [5].

Results for the additional datasets are shown in **Table** S11. HyenaDNA achieved the highest SCC score across all models.

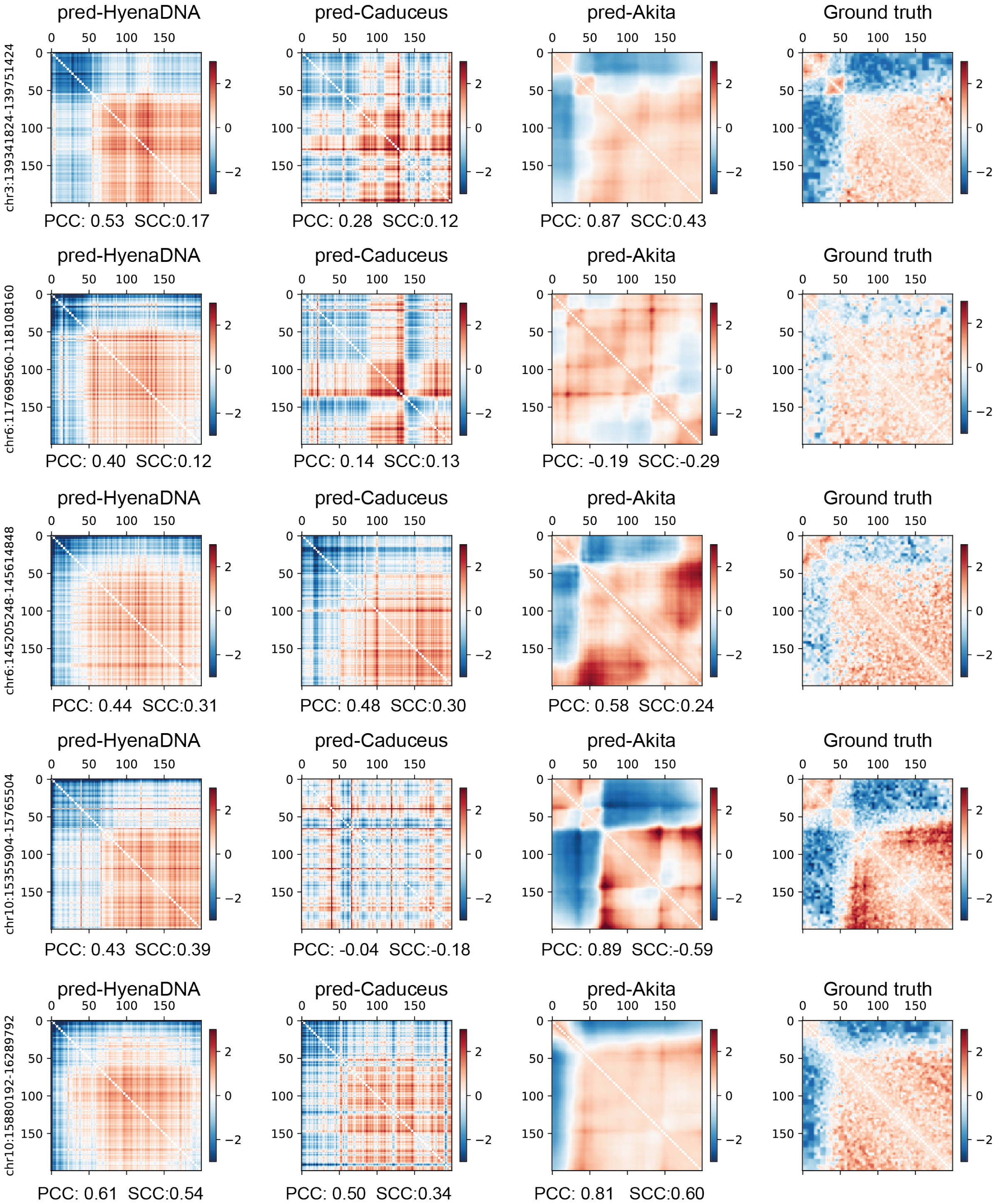


**Figure S1:** Comparisons of HyenaDNA, Caduceus (Ph), and the Expert Model (Akita) on the 2D contact map prediction task spanning 409,600 bp with a bin size of 2,048 bp. The columns show the contact map predicted by HyenaDNA, Caduceus, Akita model, and the ground truth contact map for five different genomic regions (rows). Colors represent the intensity of contact frequency between paired loci. Pearson correlation coefficient (PCC) and stratum-adjusted correlation coefficient (SCC) metrics are shown beneath each contact map to indicate prediction performance relative to the ground truth.

| Models | CMP | | | | | |
|---|---|---|---|---|---|---|
| | HFF | H1hESC | GM12878 | IMR90 | HCT116 | Avg |
| Expert Model | **0.633** | **0.653** | **0.581** | 0.595 | 0.553 | **0.603** |
| CNN | 0.098 | 0.074 | 0.082 | 0.077 | 0.028 | 0.072 |
| HyenaDNA | 0.520 | 0.517 | 0.539 | **0.635** | **0.568** | 0.556 |
| Caduceus-Ph | 0.539 | 0.524 | 0.418 | 0.473 | 0.536 | 0.498 |
| Caduceus-PS | 0.528 | 0.454 | 0.391 | 0.424 | 0.508 | 0.461 |

**Table S1:** Pearson correlation scores for the contact map prediction (CMP) task. K562, HFF, H1hESC, GM12878, IMR90, and HCT116 represent different human cell types. The highest scores are highlighted in bold. “Avg” denotes the average score across all cell types.

| Models | HFF | H1hESC | GM12878 | IMR90 | HCT116 | Avg |
|---|---|---|---|---|---|---|
| Caduceus-Ph-409600bp | 0.153 | 0.130 | 0.101 | 0.138 | 0.145 | **0.133** |
| Caduceus-Ph-307200bp | 0.066 | 0.076 | 0.051 | 0.054 | 0.149 | 0.079 |
| Caduceus-Ph-204800bp | 0.082 | 0.090 | 0.047 | 0.053 | 0.146 | 0.083 |

**Table S2:** Ablation study on context length for the contact map prediction task.

| Models | RSAP | | |
|---|---|---|---|
| | Human | Mouse | Avg |
| HyenaDNA-196k | 0.298 | 0.396 | **0.347** |
| HyenaDNA-131k | 0.268 | 0.393 | 0.331 |

**Table S3:** Ablation study on context length for the regulatory sequence activity prediction task.

| Models | ETGP |
|---|---|
| Caduceus-Ph | **0.826** |
| Caduceus-Ph w/ shuffled context | 0.418 |

**Table S4:** Ablation study on context length for the enhancer-target gene prediction task.

| Models | eQTLP | | | | | | | | | |
|---|---|---|---|---|---|---|---|---|---|---|
| | CCF | WB | Thyroid | SNSES | SSELL | MS | NT | AT | AS | Avg |
| Caduceus-Ph | 0.597 | 0.594 | 0.527 | 0.586 | 0.574 | 0.538 | 0.588 | 0.547 | 0.541 | **0.565** |
| w/ shuffled context | 0.519 | 0.515 | 0.507 | 0.533 | 0.539 | 0.521 | 0.542 | 0.507 | 0.504 | 0.521 |

**Table S5:** Ablation study on context length for the eQTL prediction task.

| Models | TISP | | | | | |
|---|---|---|---|---|---|---|
| | FC | EC | ER | GC | PC | Avg |
| Puffin-D | 0.593 | 0.331 | 0.294 | 0.249 | 0.394 | **0.372** |
| Puffin-D w/ shuffled context | 0.530 | 0.283 | 0.450 | 0.212 | 0.331 | 0.361 |

**Table S6:** Ablation study on context length for the TISP task on the validation chromosome 10.

| Models | TISP | | | | | |
|---|---|---|---|---|---|---|
| | FC | EC | ER | GC | PC | Avg |
| Puffin-D | 0.808 | 0.710 | 0.749 | 0.624 | 0.774 | **0.733** |
| Puffin-D w/ shuffled context | 0.781 | 0.699 | 0.715 | 0.604 | 0.702 | 0.700 |

**Table S7:** Ablation study on context length for the TISP task on the test chromosomes 8-9.

| Models | ETGP |
|---|---|
| Expert Model (ABC) | 0.896 |
| Expert Model (gABC) | 0.890 |
| CNN | 0.711 |
| HyenaDNA | 0.752 |
| Caduceus-Ph | **0.906** |
| Caduceus-PS | 0.895 |

**Table S8:** ETGP results on data curated from [4].

| Models | ETGP |
|---|---|
| Expert Model (ABC) | 0.824 |
| Expert Model (gAB) | **0.831** |
| CNN | 0.730 |
| HyenaDNA | 0.781 |
| Caduceus-Ph | 0.788 |
| Caduceus-PS | 0.793 |

**Table S9:** ETGP results on data curated from [3].

| Cell Type | Accession Number | Data Source |
|---|---|---|
| HAP1 | 4DNFIWGGYEW2 | 4DN |
| Hela | 4DNFI65WJKMT | 4DN |
| HepG2 | 4DNFIQ4G74OW | 4DN |
| K562 | 4DNFI2R1W3YW | 4DN |

**Table S10:** Accession number and data source of the additional datasets curated for the contact map prediction task.

| Models | CMP | | | | |
|---|---|---|---|---|---|
| | HAP1 | Hela | HepG2 | K562 | Avg |
| Expert Model | **0.196** | **0.223** | **0.198** | **0.175** | **0.198** |
| CNN | 0.018 | 0.025 | 0.021 | 0.003 | 0.017 |
| HyenaDNA | -0.062 | 0.103 | 0.094 | 0.065 | 0.049 |
| Caduceus-Ph | 0.063 | 0.168 | 0.178 | 0.004 | 0.103 |
| Caduceus-PS | 0.063 | 0.170 | 0.178 | 0.051 | 0.115 |

**Table S11:** Stratum-adjusted correlation coefficient (SCC) for the contact map prediction (CMP) task on additional datasets. HAP1, Hela, HepG2, K562 represent different human cell types. The highest scores are highlighted in bold. “Avg” denotes the average score across all cell types.
